# A method for mining condition-specific co-expressed genes in *Camellia sinensis* based on K-means clustering: A case study of “Anji Baicha” tea cultivar

**DOI:** 10.1101/2024.01.25.577317

**Authors:** Xinghai Zheng, Peng Ken Lim, Marek Mutwil, Yuefei Wang

## Abstract

As one of the world’s most important beverage crops, tea plants (*Camellia sinensis*) are renowned for their unique flavors and numerous beneficial secondary metabolites, attracting researchers to investigate the formation of tea quality. With the increasing availability of transcriptome data on tea plants in public databases, conducting large-scale co-expression analyses has become feasible to meet the demand for functional characterization of tea plant genes. However, as the multidimensional noise increases, larger-scale co-expression analyses are not always effective. Analyzing a subset of samples generated by effectively downsampling and reorganizing the global sample set often leads to more accurate results in co-expression analysis. Meanwhile, global-based co-expression analyses are more likely to overlook condition-specific gene interactions, which may be more important and worthy of exploration and research. Here, we employed the k-means clustering method to organize and classify the global samples of tea plants, resulting in clustered samples. Metadata annotations were then performed on these clustered samples to determine the “conditions” represented by each cluster. Subsequently, we conducted gene co-expression network analysis (WGCNA) separately on the global samples and the clustered samples, resulting in global modules and cluster-specific modules. Comparative analyses of global modules and cluster-specific modules have demonstrated that cluster-specific modules exhibit higher accuracy in co-expression analysis. To measure the degree of condition specificity of genes within condition-specific clusters, we introduced the correlation difference value (CDV). By incorporating the CDV into co-expression analyses, we can assess the condition specificity of genes. This approach proved instrumental in identifying a PPR-type RNA editing factor gene (CWM1) that specifically functions during the bud-prealbinism stage of the *Camellia sinensis* cultivar “Anji Baicha”. We hypothesize that this gene may be upregulated and play a role in inhibiting chloroplast development, ultimately resulting in albino phenotypes in “Anji Baicha”.

## Introduction

As one of the most popular non-alcoholic beverages worldwide, tea contains a wide range of secondary metabolites beneficial to human health, such as polyphenols, alkaloids, and amino acids (Wang et al., 2022). As such, the tea plant (*Camellia sinensis*) possesses a diverse range of germplasm resources (Chen et al., 2007). Different cultivars of *Camellia sinensis* are each prized for certain desirable qualities in their own right and exhibit significant differences in plant morphology, leaf characteristics, growth habits, adaptability, and secondary metabolites (Chen et al., 2012; Zhao et al., 2022). Consequently, said tea cultivars have garnered much research interest in the post-genomic era to understand and improve tea traits. For example, temperature-sensitive albino and light-sensitive yellowing varieties have attracted much attention due to their unique leaf color and secondary metabolite content (Zhang et al., 2020). Taking the amino acid-rich “Anji Baicha” variety as an example, its new shoots are highly sensitive to cold temperatures and undergo a color transformation as the temperature gradually warms up in early spring. In early spring, the tender leaves of “Anji Baicha” appear white and gradually transition to light green leaves as the temperature rises, returning to normal green color within approximately two weeks (Cheng et al., 1999a; Li, 2002). Meanwhile, the amino acid content in dry tea made from the shoots of “Anji Baicha” during the white stage can reach up to 6%, three times higher than that of ordinary cultivars (2%) (Li et al., 1996; Xiong et al., 2013).

With an increasing number of studies on the epigenetic variations and compositional changes of secondary metabolites in tea plants under different experimental conditions (Zhao et al., 2022; Wang et al., 2022; Liao et al., 2022), the omics dataset of *Camellia sinensis* has also become increasingly extensive. This has led to the use of systems biology approaches on sequencing data hosted on public databases (Tai et al., 2018; Xia et al., 2020; Zhao et al., 2021), such as gene co-expression analysis, becoming a trend in analyzing omics data of *Camellia sinensis*, providing tea researchers with a more macroscopic and comprehensive perspective. Researchers have further downloaded large-scale transcriptome data of tea plants and created a more systematic and comprehensive co-expression database TeaCoN (http://teacon.wchoda.com) (Zhang et al., 2020).

Although the large sample size of publicly-derived *Camellia sinensis* transcriptomic data improves the statistical significance of relationships between genes and increases the reliability of inferring gene correlations, indiscriminately combining multiple samples may not be universally beneficial (He and Maslov, 2016). As datasets become larger and more diverse, the derived coexpression networks become less informative due to increased multidimensional noise (Liesecke et al., 2019). One way to improve the utility of the network is downsampling. Downsampling subdivides samples either by manually grouping them based on experimental conditions or by using automated methods such as k-means clustering (Feltus et al., 2013; Gibson et al., 2013; Xiao et al., 2014). However, manual grouping often lacks sufficient sample description to accurately classify them, so automated methods like k-means clustering are more effective (Feltus et al., 2013).

Furthermore, co-expression networks at a large scale of samples may miss specific gene interactions formed under particular conditions (Fuente et al., 2010). Increasing evidence suggests that different gene networks operate in different biological contexts (Roguev et al., 2008; Bandyopadhyay et al., 2010). Therefore, it becomes increasingly important to compare and contrast coexpression networks under specific conditions (Choi et al., 2005; Ideker et al., 2012; Amar et al., 2013). Experimental results demonstrate that over one-third of genetic interactions are condition-specific (Guénolé et al., 2013). Several studies have also shown that the patterns of gene coexpression vary under different conditions (Southworth et al., 2009; Hudson et al., 2009; Anglani et al., 2014). Hence, when conducting coexpression analysis on large-scale samples, incorporating sample auto-classification and mining condition-specific coexpressed genes can enhance the accuracy and informativeness of co-expression network analysis.

In this study, all *Camellia sinensis* samples downloaded from NCBI were subjected to k-means clustering to obtain seven clusters representing different “conditions” (experimental treatments, tissues, and cultivars). Cluster metadata annotations were obtained through sample metadata annotation. Then, weighted gene co-expression network analysis (WGCNA) was performed on the expression profiles of both the global samples and the cluster samples to obtain their respective co-expression modules. Subsequently, the correlation difference value (CDV) was proposed to measure the degree of condition specificity of genes within condition-specific clusters. By comparing between clusters and within clusters, highly condition-specific clusters and biological functions were identified. By incorporating the correlation difference value (CDV) into coexpression networks and visualizing it, condition-specific genes and conserved genes can be distinguished, providing more information for the selection of key genes. Overall, this study aims to improve gene co-expression analysis methods for large-scale transcriptomic data of tea plants by performing condition-specific analysis and providing a more accurate understanding of the relationships between gene expression patterns and phenotypic traits.

## Methods

### Data sources and sample metadata annotation

By searching and filtering using the keyword “Camellia sinensis” in the NCBI SRA database, a total of 769 RNA-Seq raw reads were obtained. Initial annotations of these RNA-Seq raw data were performed using the metadata fields in the NCBI SRA database, including cultivar, plant tissue, and experimental treatments. Subsequently, the corresponding original papers for each RNA-Seq data were searched to retrieve annotation information (Table S1).

For the control group, samples in each experiment or samples directly collected without any treatment were uniformly labeled as “no treatment” in the experimental treatment column. The samples with missing annotations in the metadata fields of the NCBI SRA database and could not be found in the retrieved original papers were labeled as “missing”.

### Expression quantification and gene functional annotation

769 RNA-seq samples were processed using fastp tool (Chen et al., 2018) to obtain high-quality clean data by removing adapter sequences and low-quality reads using default parameters. Coding sequences (CDS) annotations of the “Shuchazao” *Camellia sinesis* cultivar (http://tpia.teaplant.org) (Xia et al., 2019) were used as pseudoalignment reference, the processed reads were then used to quantify the gene expression, in transcripts per million (TPM) values, for all RNA-seq samples using Kallisto (Bray et al., 2016) (Table S2). Additionally, the percentage of reads pseudoaligned to the CDS file in the clean data was calculated as the pseudoaligned reads percentage, which was used to assess sample quality (Figure S1; Table S1) as described in (Tan et al., 2020; Hew et al., 2020).

The CDS annotations of the tea plant cultivar “Shuchazao” were subjected to gene functional annotation using the Mercator v4 2.0 (Lohse et al., 2014) (Table S3).

### K-means clustering and cluster metadata annotation

Firstly, a gene expression profile was constructed using transcripts per million (TPM) values of 22,842 genes from 769 RNA-seq samples. Then, the StandardScaler tool from the sklearn.preprocessing package was used to standardize TPM values of the gene expression profile within samples. Next, the KMeans tool from the sklearn.cluster package was used for K-means clustering on all RNA-seq samples with a random seed set to 1024 (np.random.seed(1024)) and a target number of clusters set to 7 (n_clusters=7) (Tavazoie et al., 1999). The selection of 7 as the value of k in K-means clustering is based on pre-experimental data, where 7 resulted in a better “conditional” classification of the samples (Table S1).

Then, the PCA tool from the sklearn.decomposition package was applied to the standardized gene expression profile to perform dimensionality reduction, retaining the top principal components PC1 to PC3, which explains the majority of the variability in the expression data of the samples. Finally, the samples were visualized in the PC1 to PC3 space to explore potential clustering structures and similarities among the samples, as described by (Clayman et al., 2020).

To annotate the 7 clusters obtained from K-means clustering, the hypergeom tool from the scipy.stats package was used to perform a hypergeometric test between each sample in each cluster and the samples associated with each cultivar term (Hahne et al., 2008). Then, the fdrcorrection tool from the statsmodels.stats.multitest package was used to correct the p-values of all cultivar terms corresponding to each cluster, obtaining the false discovery rate (FDR) values (Benjamini and Hochberg, 1995). Cultivar terms with FDR values less than or equal to 0.05 were selected as metadata annotations for the samples in that cluster. The same approach was applied to obtain metadata annotations for the tissue terms and experimental treatment terms (Figure S2; Table S4).

### Weighted gene co-expression network analysis (WGCNA)

The global expression profile and cluster expression profiles comprise the expression levels of 22,842 genes from global samples and samples from different clusters (cluster samples), respectively.

R package WGCNA was then employed to construct a co-expression network (Langfelder et al., 2008). When constructing the co-expression network, an appropriate soft threshold was selected based on the scale-free topology fit index and the mean connectivity. By analyzing the scale-free topology fit index plot, the first point that reached a value above 0.9 was identified as the soft threshold (Figure S3).

Next, WGCNA was performed following the guidelines provided in the tutorial (https://horvath.genetics.ucla.edu/html/CoexpressionNetwork/Rpackages/WG CNA/Tutorials/) (Zhang and Horvath, 2005), where employing a stepwise approach to divide the modules based on the obtained soft threshold and utilized the DynamicTreecut package to cluster the modules.

The eigengenes of each module were calculated based on the expression profiles and module color codes (Figure S4). Eigengenes represent the main expression patterns of each module and can be used to describe the overall expression patterns of the module. Then, hierarchical clustering (average linkage) was applied to cluster the module eigengenes. A merging height threshold of 0.25 was set, corresponding to a correlation threshold of 0.75, and called the mergeCloseModules function for automatic module merging. The merged modules were assigned new color codes, which served as the final module color codes (Figure S5). For easy reference, the color modules were mapped to alphabetical letters (Table S5).

In this step, the co-expression modules obtained from the global expression profile are referred to as “global modules”, while the co-expression modules obtained from the cluster expression profiles are referred to as “cluster-specific modules”.

### Calculation of clustering similarity and gene condition specificity

The Fowlkes-Mallows score (FMS) and adjusted mutual information score (AMIS), which is used to analyze the similarity of co-expression modules in WGCNA under different samples, was calculated using the fowlkes_mallows_score and adjusted_mutual_info_score tool from the sklearn.metrics package (Fowlkes and Mallows, 1983; Amelio and Pizzuti, 2017).

Then, for each cluster, the two kinds of gene-module consistency coefficient (GMC) were calculated for each gene - the GMC of the gene in the cluster-specific module and the GMC of the gene in the global module based on corresponding cluster samples. The GMC of a gene is defined as the Pearson correlation coefficient (PCC) between the gene’s expression profile and the average expression profile of the co-expression module it belongs to (Cohen et al., 2009). This study used the GMC of genes to investigate the consistency of gene expression patterns within co-expression modules.

Furthermore, for each cluster, the correlation difference value (CDV) of each gene was calculated by subtracting the GMC of the gene in the cluster-specific module from the GMC of the gene in the global module. In this study, CDV was used to measure the condition-specificity of each gene.

### Module functional annotation and module correlation analysis

For module correlation analysis, the hypergeometric test (Hahne et al., 2008) was performed using the hypergeom tool from the scipy.stats package for each gene in the global expression profile and each gene in the cluster expression profile of each module. Then, the p-value of each global module corresponding to each cluster-specific module was corrected using the fdrcorrection tool from the statsmodels.stats.multitest package to obtain the false discovery rate (FDR) values (Benjamini and Hochberg, 1995). Module pairs with FDR values less than or equal to 0.01 in the “global-cluster” relationship were considered to have significant module correlations (Figure S6; Table S6).

To annotate the functions of modules, the hypergeometric tests (Hahne et al., 2008) were performed using the hypergeom tool from the scipy.stats package for each gene in the module and each gene included in the mapman entries (only mapman entries with detailed classification and containing more than 100 genes are selected). Then, the p-values of all mapman entries corresponding to each module are corrected using the fdrcorrection tool from the statsmodels.stats.multitest package to obtain FDR values (Benjamini and Hochberg, 1995). Mapman entries with FDR values less than or equal to 0.05 are considered functional annotations for the genes in that module (Figure S7; Table S7).

### Target gene set filtering and co-expression network construction

Based on module correlation and module functional annotation, the gene set associated with the metadata annotation of the cluster was selected. This gene set includes genes that are enriched in the target biological functions identified within the target module. Then, the Pearson correlation coefficient (PCC) values were calculated between all genes in the gene set using the cluster expression profile (Cohen et al., 2009). To construct the co-expression network, only gene pairs with a PCC greater than or equal to 0.7 were considered.

The constructed gene interaction networks were imported into Cytoscape software (Shannon et al., 2003) for visualization and analysis. In Cytoscape, the betweenness centrality of each gene was calculated (Table S8). Finally, the network was further customized with layout, labeling, and color coding to provide a clearer representation of the interactions between genes to understand the structure and function of the gene network and to reveal important associations and regulatory mechanisms in biological processes.

## Results

### Metadata annotation of RNA-Seq samples exhibits an imbalanced distribution characteristic

During the metadata annotation of 769 RNA-seq samples of *Camellia sinensis*, we observed that there was an imbalance in the sample distribution across each metadata category, including cultivars, tissues, and treatments (Figure 1; Table S1). Specifically, certain categories within tissues and treatments have a higher number of samples compared to others. For example, in the 769 *Camellia sinensis* RNA-seq samples, the “leaf” samples accounted for 53.7% of the total, while the “no treatment” samples in the experimental treatments category accounted for 46.2%, far exceeding the numbers of other categories (Figure 1). Regarding cultivars, we saw a relatively balanced representation across different categories, but some cultivars have a higher proportion. For instance, “Shuchazao” accounts for 17.6%, “Longjing 43” accounts for 17.0%, and “Fuding Dabaicha” accounts for 11.4%.

**Figure 1.**
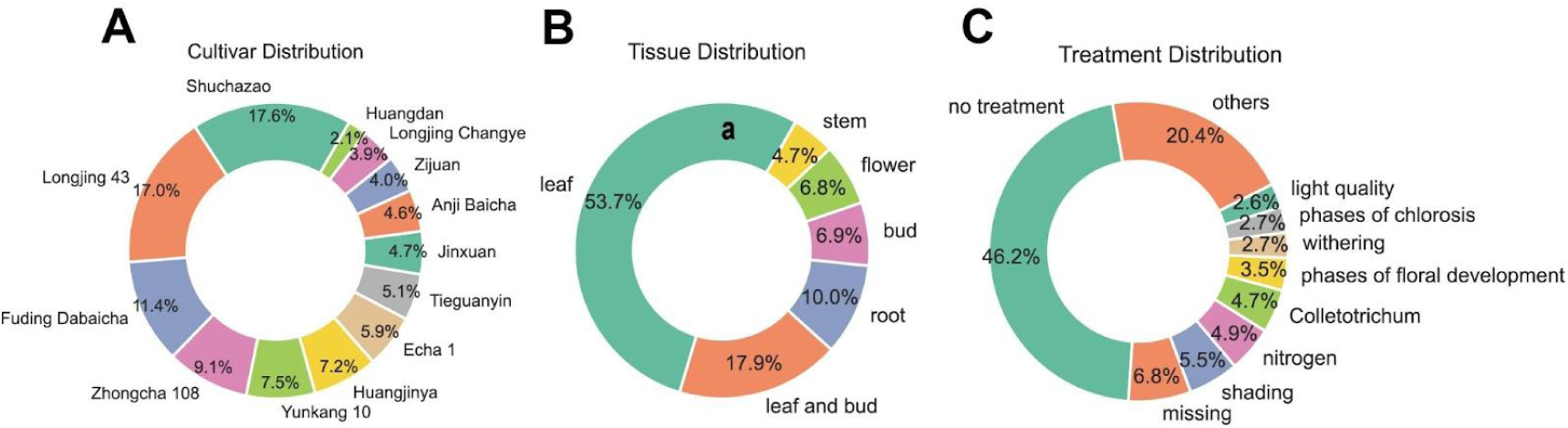
Analysis of metadata for RNA-seq samples of *Camellia sinensis*. (A) Cultivar. (B) Tissue. (C) Experimental treatments.

### Metadata annotation of K-means clustering clusters represents specific “conditions”

K-means clustering is used to organize and classify the globally imbalanced samples in the metadata term. The metadata annotations of the clustered samples are then used as the “conditions” representing the specificity of the cluster samples.

After analyzing 769 *Camellia sinensis* RNA-seq samples using the K-means clustering algorithm, seven clusters were obtained. Among them, Cluster 7, Cluster 6, and Cluster 5 accounted for 21.8% (168 samples), 21.6% (166 samples), and 21.5% (165 samples) of the total samples, respectively, while the smallest cluster, Cluster 2, accounted for only 5.3% (41 samples) (Figure 2A) (Table 1).

**Figure 2.**
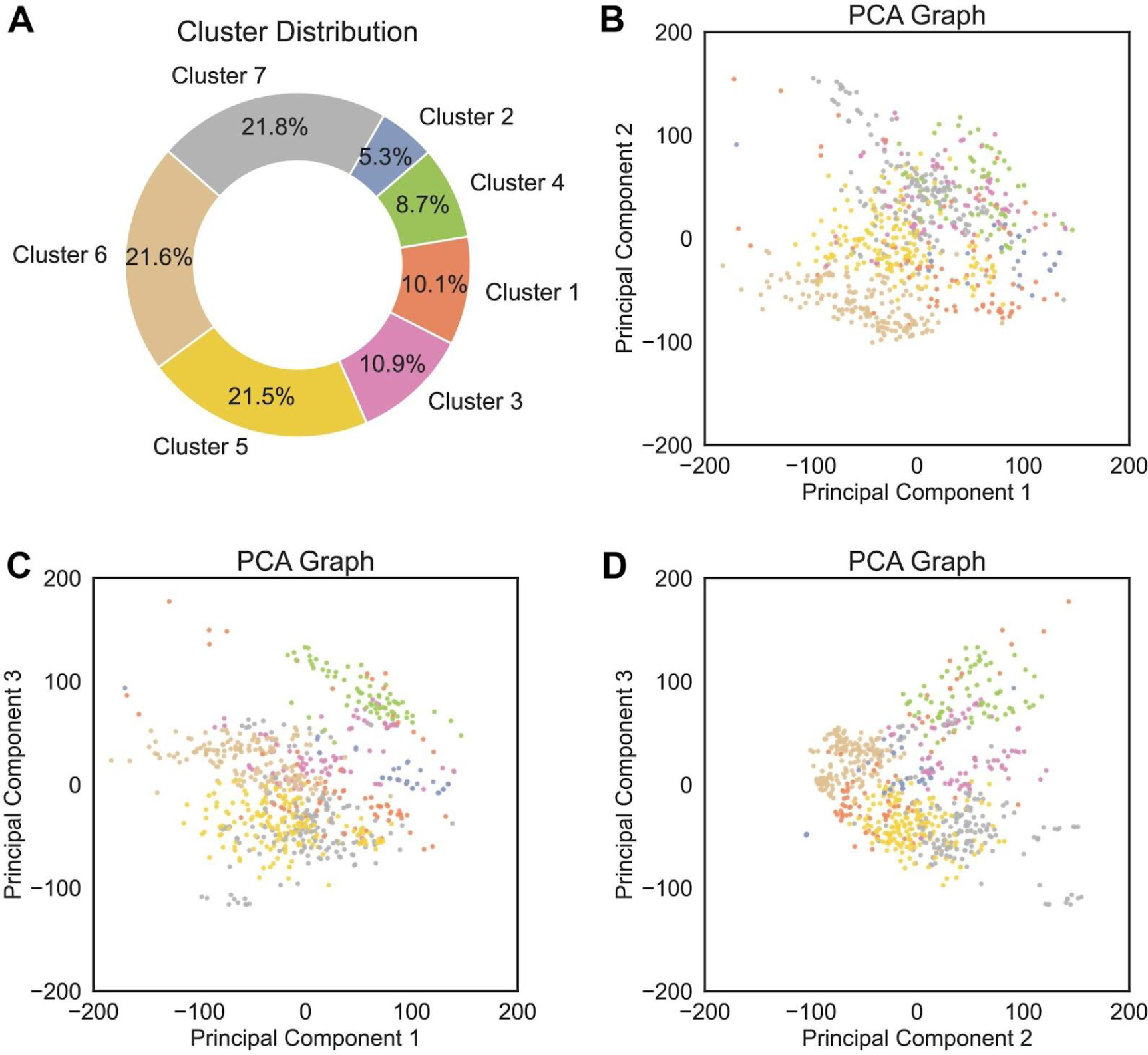
K-means clustering and principal component analysis (PCA) for *Camellia sinensis* RNA-seq samples. (A) Pie chart showing the proportion of k-means clusters. (B-D) Scatter plots of PCA show the spatial distribution of all *Camellia sinensis* RNA-seq samples on PC1 to PC3. Different clusters are distinguished using different colors, while the same cluster remains consistent across a-d.

**Table 1.** Metadata annotation table of k-means clusters.

| Clusters | Cultivar | Tissue | Treatment | # Modules | # Samples |
| --- | --- | --- | --- | --- | --- |
| Cluster 1 | Anji Baicha | leaf and bud | phases of chlorosis, continuous cold stress, gibberellin, melatonin | 34 | 78 |
| Cluster 2 | Longjing Changye, Shuchazao | flower | phases of floral development | 26 | 41 |
| Cluster 3 | Echa 1, Longjing 43 | bud, stem | NAA, Colletotrichum, withering, missing | 34 | 84 |
| Cluster 4 | Fuding Dabaicha, Zhongcha 108 | root | selenite, nitrogen, As, Cd, Na<sub>2</sub>SeO<sub>3</sub> | 28 | 67 |
| Cluster 5 | Huangjinya, Tieguanyin, Zhongcha 108, Zijuan | leaf | no treatment, Colletotrichum, fluoride, salicylic acid, sucrose | 48 | 165 |
| Cluster 6 | Shuchazao | leaf and bud, bud | no treatment, shading, drought | 35 | 166 |
| Cluster 7 | Huangjinya, Jinxuan | leaf, stem | freezing, selenite, insect infestation, light quality, mechanical damaged, UV-B, chilling, heat stress | 45 | 168 |

By performing principal component analysis (PCA) on the transcriptome data to reduce the dimensionality of the genes, the spatial distribution of these samples in PC1 to PC3 was observed (Figure 2B-D).

By conducting enrichment analysis on the metadata of k-means clusters (Figure S2), metadata terms related to cultivars, tissues, and treatments were annotated to each k-means cluster, facilitating a better understanding of the characteristics and functions of *Camellia sinensis* RNA-seq samples represented by each k-means cluster (Table 1). For example, Cluster 1 mainly includes leaves and buds of the “Anji Baicha” cultivar, with experimental treatments focused on different stages of leaf whitening, sustained low-temperature stress, gibberellins, and melatonin, among others. Such annotations are also called “conditions” represented by Cluster 1.

### Comparative analysis reveals differences between the global and cluster-specific co-expression modules

Weighted gene co-expression network analysis was used to obtain global modules and cluster-specific modules from global samples and cluster samples. 56 co-expression modules were obtained based on the global expression profile, indicating the presence of complex and diverse co-expression relationships among genes (Table S5). Different numbers of co-expression modules were obtained based on the cluster expression profiles. Specifically, 34, 26, 34, 28, 48, 35, and 45 co-expression modules were obtained based on Cluster 1 to Cluster 7 expression profiles, respectively (Table 1). The varying numbers of cluster-specific co-expression modules reflect changes in gene expression patterns under different cultivars, tissues, and experimental treatments.

Three perspectives of analysis were performed to elucidate the extent of differences in co-expression modules obtained from global samples and cluster samples: similarity analysis of module genes, internal consistency analysis of module expression profiles, and enrichment analysis of module biological functions.

To investigate the similarity of co-expression modules obtained from global samples and cluster samples in WGCNA, the Fowlkes-Mallows score (FMS) was used as a metric. FMS is commonly used to compare the similarity of clusters or co-expression modules obtained from different samples or conditions. The FMS score ranges from 0 to 1, where a value closer to 1 indicates a higher similarity between the two data sets. Conversely, when FMS approaches 0, it indicates a low consistency between the two datasets. We observed that the FMS between the global module and cluster-specific modules ranges from 0.1 to 0.2 (Figure 3A), indicating a generally low similarity between the global module and cluster-specific modules. Specifically, the similarity between the global module and the cluster-specific modules of Cluster 1, Cluster 3, and Cluster 6 is low. Additionally, the cluster-specific module of Cluster 1 shows low similarity with the majority of other clusters’ specific modules. This suggests a higher level of uniqueness for Cluster 1, suggesting that Anji Baicha cultivar might employ a transcriptional program different from the other cultivars.

**Figure 3.**
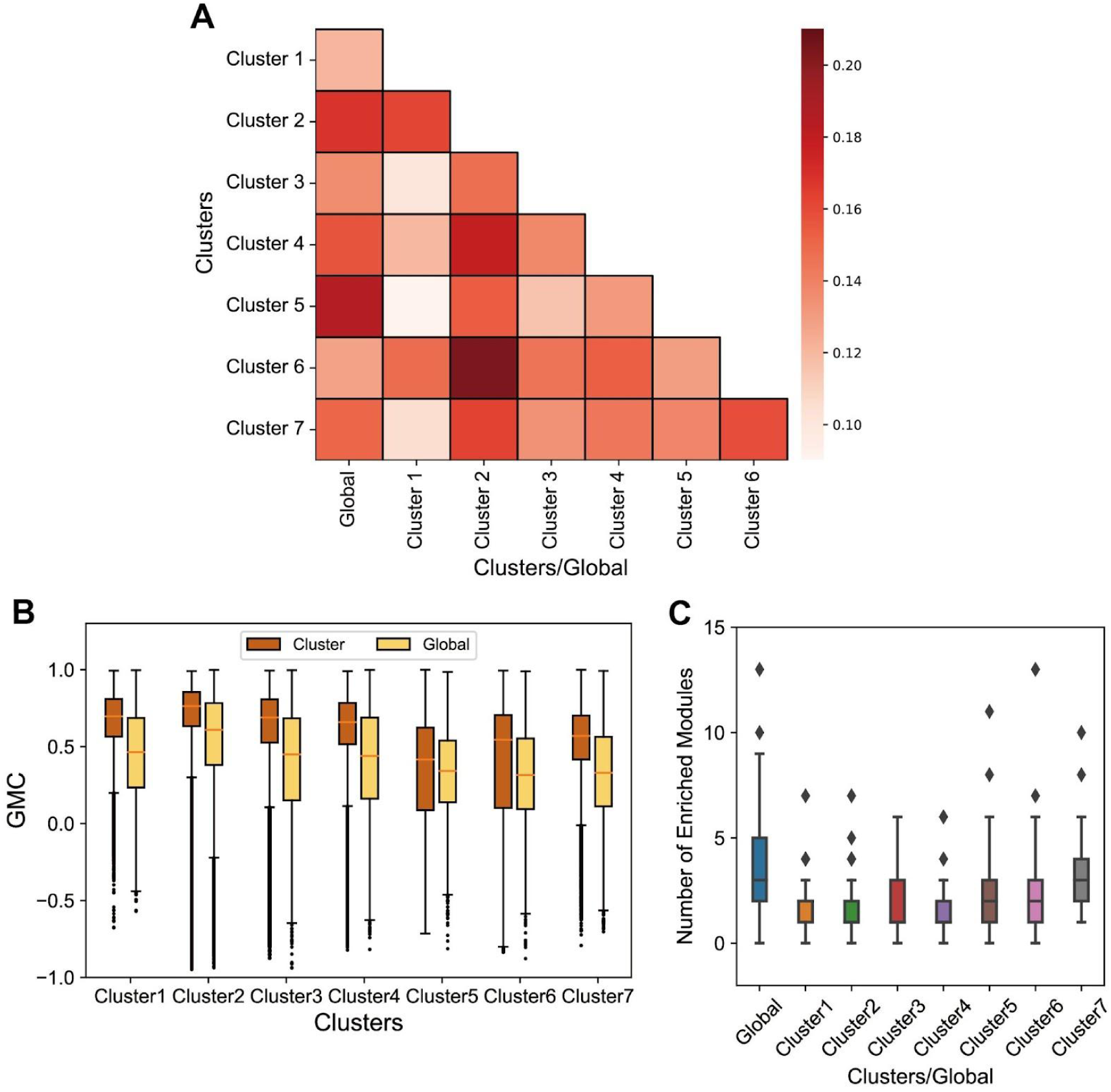
Comparative analysis of global and cluster-specific co-expression modules. (A) Similarity analysis of global and cluster-specific co-expression modules. The intensity of colors in the heatmap represents the magnitude of the Fowlkes-Mallows score (FMS). (B) Comparison of the gene-module consistency coefficient (GMC) of all genes between the global module and the cluster-specific module for each cluster. (C) Comparison of the module enrichment degree (MED) among cluster-specific modules and the global module.

We used the gene-module consistency coefficient (GMC) to assess the similarity between gene expression profiles within a module and the average expression profile of the module. The GMC is essentially the correlation coefficient between the gene expression profile within the module and the average expression profile of the module. It ranges from −1 to 1, where a value greater than 0 indicates a positive correlation and a value less than 0 indicates a negative correlation. As expected, genes from the same module tend to have a GMC score >0, as genes in the same module should be correlated (Figure 3B). However, the median GMC of genes in the cluster-specific modules tends to be higher than the GMC of genes in the global module (Figure 3B), with exception of Cluster 5, which is the cluster most similar to the global module (Figure 3A). This shows that choosing biologically related samples increases correlations between genes and might improve the biological inferences.

Lastly, we used module enrichment degree (MED) to examine the distribution of various biological functions within the global module and specific modules. MED refers to the number of modules in which a biological function is enriched.

A higher MED value indicates that the biological function is more dispersed among modules, while a lower MED value indicates that the biological function is more concentrated. We observed that, compared to the global module, the biological function is more concentrated in the cluster-specific modules of Cluster 1, Cluster 2, and Cluster 4 (Figure 3C). A higher MED value indicates that a biological function is enriched in more modules, while a lower MED value indicates that a biological function is enriched in fewer modules. We found that compared to global modules, the median MED value is significantly lower in cluster-specific modules, which is reasonable. This is because the partitioning of global modules is based on global samples. Global samples have diverse treatments and conditions, which helps activate more biological functional modules in tea plants.

### Selecting the condition-specific co-expression modules and biological functions from two perspectives

In traditional WGCNA, after obtaining co-expression modules, there is often a biological functional annotation of the modules. However, here, we not only annotate the modules with functional information, but also calculate the conditional specificity of each module to each function. In this study, correlation difference value (CDV) is proposed as a measure of gene condition specificity. CDV is calculated as the difference between the Gene Module Consistency Coefficient (GMC) of a gene in the cluster-specific module and its GMC in the global module. CDV values range from −2 to 2. A CDV value closer to 2 indicates a higher level of gene condition specificity, while a value closer to 0 indicates a higher level of conservation, as the average expression of the gene is more similar to the expression profile of the global module.

To explain the biological function of condition-specific genes, we analyzed a series of CDV thresholds ranging from 0 to 0.8. For each threshold, genes with CDV values higher than the threshold were considered cluster-specific, while genes with CDV values lower than the threshold were considered conserved. For each threshold, we calculated two values: the similarity between the global module and the cluster-specific module after removing genes with values higher than the threshold (red line), and the similarity between the global module and the cluster-specific module after removing genes with values lower than the threshold (blue line). Therefore, the red line can be understood as follows: when the threshold is close to 0, there is a high similarity between the global module and the cluster-specific module. However, as the threshold increases from 0 to 0.8, genes with higher CDV values are included in both the global module and the cluster-specific module, resulting in a decrease in the similarity between them (Figure 4A) (Figure S8). In other words, genes with higher CDV values lead to lower similarity between the global module and the cluster-specific module, indicating higher condition specificity. On the other hand, the blue line indicates that as the threshold decreases from 0.8 to 0, genes with lower CDV values are included in both the global module and the cluster-specific module. Genes with lower CDV values have minimal impact on the similarity between the global module and the cluster-specific module.

**Figure 4.**
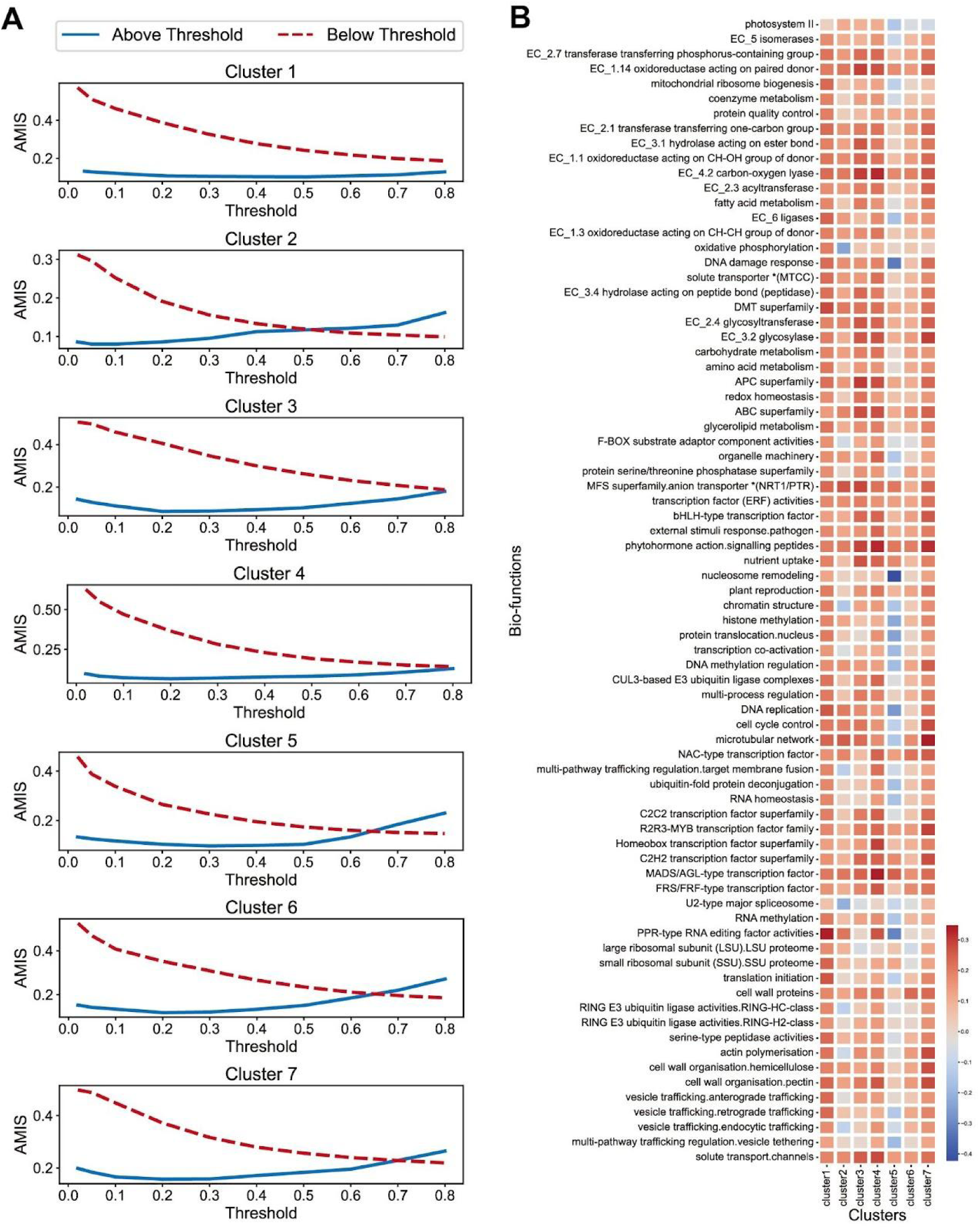
Relationship between Correlation difference value (CDV) and condition specificity, and average CDV of different biological functions in different clusters. (A) The impact of genes with different Correlation difference value (CDV) on the similarity of global modules and cluster-specific modules. (B) Correlation difference value (CDV) heatmap for each bio-function in each cluster. The color of the cells represents the average CDV.

Subsequently, the average CDV of genes with different biological functions in the seven clusters revealed that Cluster 5, which was found to be most similar to the global module (Figure 3), has a higher proportion of genes with low average CDV values associated with various biological functions (Figure 4B). Conversely, Cluster 1, which was found to be least similar to the global module, has a higher proportion of genes with high average CDV values associated with specific biological functions. This further underscores the relationship between CDV and condition specificity.

As discovered in the previous section, Cluster1-specific module is the least similar to the global module and is most likely to uncover condition-specific modules and biological functions. We analyzed Cluster 1 from two perspectives: gene condition specificity and biological function enrichment. By combining the average CDV (correlation difference value) heatmap, with the specific modules of Cluster 1 as the x-axis, and biological functions as the y-axis, and the results of significant biological function enrichment, we observed that the biological function with the highest average CDV is “PPR-type RNA editing factor activities”. This biological function is enriched in three WGCNA co-expression modules: “grey”, “yellow4”, and “lightsteelblue1” (Figure 5A). Among them, the “yellow4” and “lightsteelblue1” co-expression modules have relatively high average CDV. It can also be observed that both “yellow4” and “lightsteelblue1” are correlated with the global module “cyan”, with “yellow4” showing higher correlation and more enriched biological functions (Figure 5B). This suggests that RNA editing factors that could modify chloroplastic RNAs (Hammani et al., 2009; Cui et al., 2019), are responsible for the albino phenotype of “Anji Baicha” cultivar.

**Figure 5.**
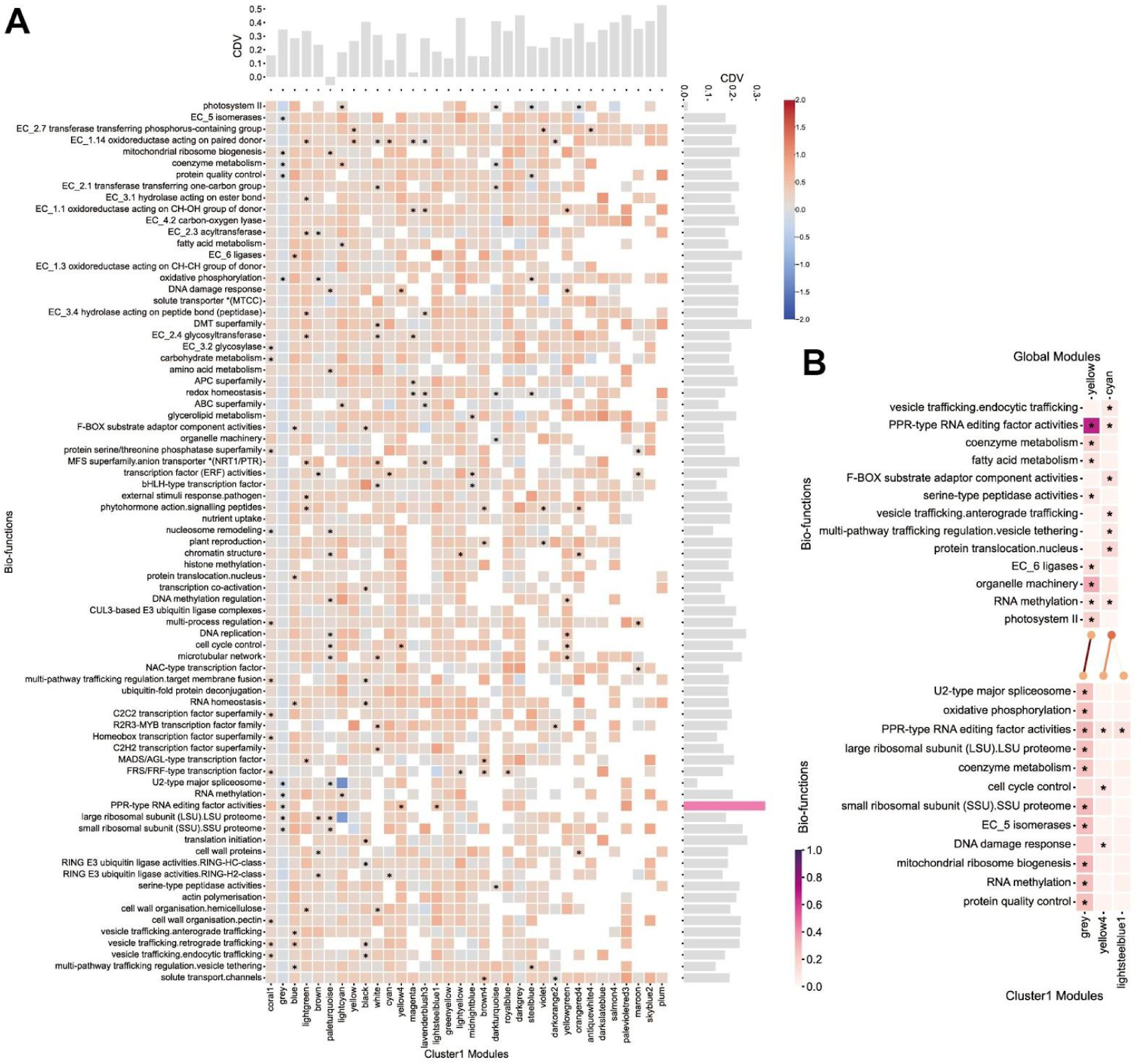
Comparative analysis of the cluster1-specific module and the global module from two perspectives: gene-module consistency coefficient (GMC) and functional enrichment. (A) Correlation difference value (CDV) heatmap and functional enrichment heatmap corresponding to various biological functions for each co-expression module in Cluster 1. Cells marked with asterisks (*) indicate significant enrichment, and the color of the cells represents the average CDV. (B) Comparison analysis of functional enrichment heatmaps related to “PPR-type RNA editing factor activities” between the global module and the cluster-specific module. Cells marked with asterisks (*) indicate significant enrichment, and the color of the cells represents the recall. The bipartite graph in the central region of the figure 5B reveals the relationship between the cluster-specific module and the global module. Nodes represent modules, and the color intensity of the nodes represents their degree. The presence or absence of connecting lines between nodes distinguishes significant or non-significant associations, and the color intensity of the connecting lines represents the Jaccard index. The size of the modules decreases from left to right.

### Combining condition specificity and co-expression network analysis reveals an important PPR-type RNA editing factor functions during the bud-prealbinism stage

In the specific condition Cluster 1 “yellow4” module, we found that the biological function “PPR-type RNA editing factor activities” exhibited the highest condition specificity. To explore this network and identify condition-specific co-expressed genes that could potentially explain the specificity of Cluster 1 under certain conditions, we extracted genes associated with several enriched biological functions in the “yellow4” module to construct a co-expression network. The aim was to examine the importance of a gene in the co-expression network from two aspects - betweenness centrality (the importance of the responsive gene in the network/biological function) and condition specificity (the importance of the responsive gene for the “condition”). This is done in order to screen out genes that are more potentially valuable for research under a given “condition”. We represented each gene with a different color based on its correlation difference value (CDV). The gene with the highest CDV is CSS0047732.1, while the gene with the lowest CDV is CSS0045541.1 (Figure 6A). This indicates that gene CSS0047732.1 exhibits a high degree of condition specificity, while gene CSS0045541.1 is more conserved.

**Figure 6.**
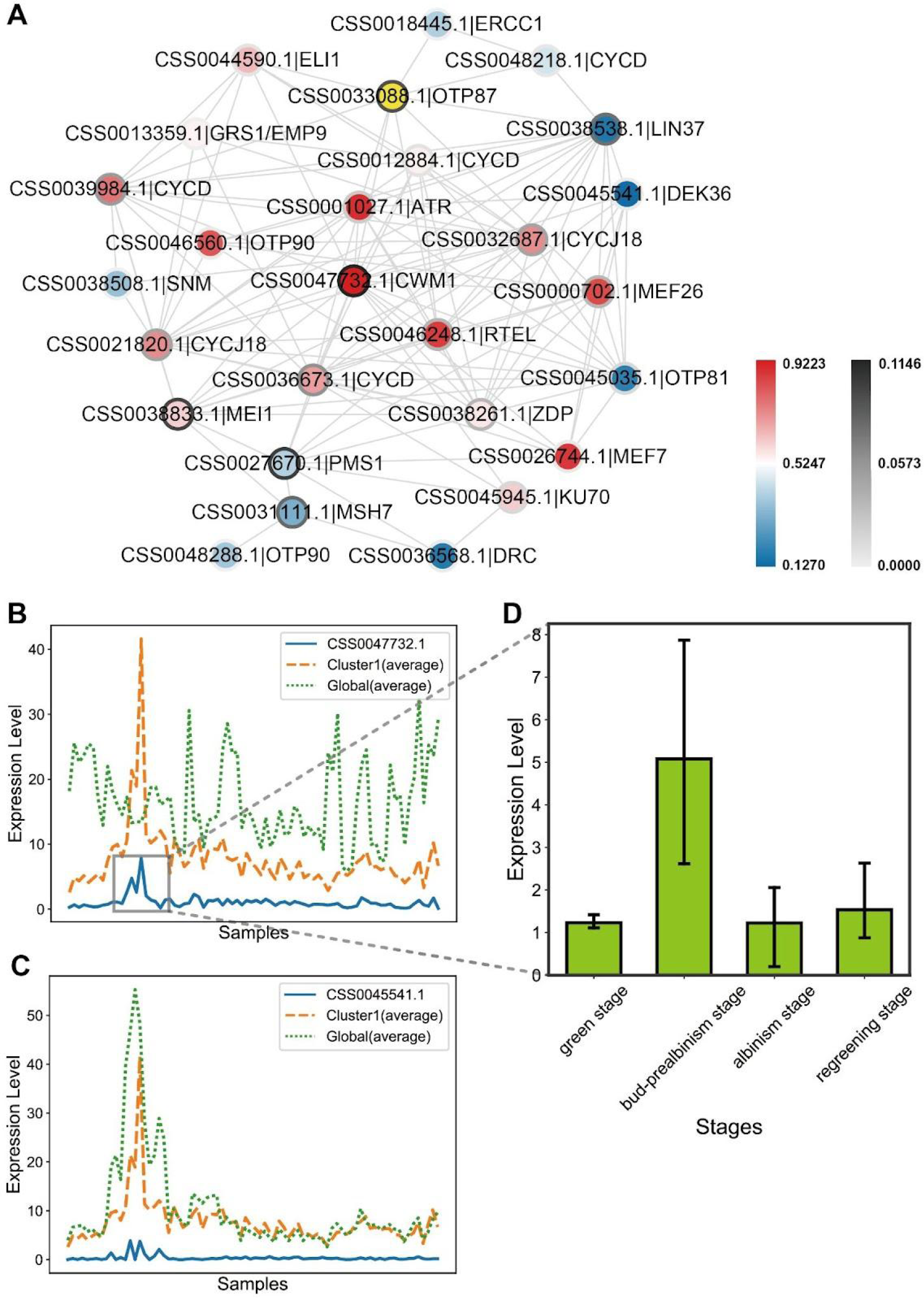
Gene co-expression network and comparison analysis of expression profiles. (A) Co-expression network of genes related to the biological function “PPR-type RNA editing factor activities” in the yellow4 module of Cluster 1. The color variation of the nodes represents the level of correlation difference value (CDV), and the color intensity of the node borders represents the betweenness centrality. (B) Comparison of the expression profile of gene CSS0047732.1 with the Cluster 1 module average expression profile and the Global module average expression profile. (C) Comparison of the expression profile of gene CSS0045541.1 with the Cluster 1 module average expression profile and the Global module average expression profile. (D) Expression levels of the CSS0047732.1 gene at different stages of leaf whitening in “Anji Baicha” during early spring.

Subsequently, by comparing the expression profiles of gene CSS0047732.1 and gene CSS0045541.1 with the average expression profiles of their respective cluster-specific module and the global module, we observed that the expression pattern of gene CSS0047732.1 is similar to the average expression profile of its cluster-specific module, while it differs significantly from the average expression profile of the global module (Figure 6B). On the other hand, gene CSS0045541.1 exhibits a different pattern. Due to the similarity between its cluster-specific module and the global module’s average expression profiles, it is challenging to determine which one its expression profile is closer to (Figure 6C). The expression profiles further validate the previous observations, with genes having higher CDV values indicating a higher degree of condition specificity, while genes with CDV values close to 0 are more conserved.

To assess the importance of each gene in the co-expression network, we added the attribute of betweenness centrality to each node. Betweenness centrality measures the importance and intermediacy of a node in the network (Table S8). A gene with high betweenness centrality indicates that it serves as a crucial bridge in the network, potentially transmitting signals or regulating information flow between different functional modules. Gene CSS0047732.1 has the highest CDV and betweenness centrality simultaneously, indicating that it not only exhibits a high degree of condition specificity but also plays a crucial role in the co-expression network (Figure 6A).

The gene CSS0047732.1 has been annotated by Mercator as a “PPR-type RNA editing factor (CWM1)”. Our investigation revealed that CSS0047732.1 exhibits significant upregulation during the early spring bud-prealbinism stage of “Anji Baicha” and downregulation during the albinism stage, indicating its functional relevance in the bud-prealbinism stage (Figure 6D). CWM1, a PPR-type RNA editing factor, exhibits high expression in the bud leaves of “Anji Baicha” during the bud-prealbinism stage, was found its active participation in the regulatory mechanisms of this specific developmental phase.

## Discussion

Tea plant (*Camellia sinensis*), being one of the world’s most important beverage crops, is known for its numerous secondary metabolites that contribute to the tea quality and health benefits. In order to characterize the biological functions of genes in tea plants, previous research has utilized a large-scale SRA data downloaded from NCBI to construct a gene co-expression network database known as TeaCoN (http://teacon.wchoda.com) (Zhang et al., 2020).

However, when conducting co-expression analysis, more samples does not necessarily mean better results (He and Maslov, 2016). Researchers analyzed a dataset of *Escherichia coli* microarray data and found that subsets of the dataset performed better in inferring transcriptional regulatory networks (Cosgrove et al., 2010). The poor performance of the global network was attributed to increased multidimensional noise (Liesecke et al., 2019). However, this issue can be mitigated by determining the optimal number of effective samples, for example, through a downsampling method that automatically groups samples using k-means clustering (Feltus et al., 2013; Gibson et al., 2013).

In this study, we observed that the metadata entries (experimental treatments, tissues, and cultivars) of the SRA samples downloaded from NCBI were imbalanced. This phenomenon has also been observed in other large-scale co-expression analysis studies (Zhang et al., 2020). The imbalanced sampling of global samples makes it difficult to represent specific research questions that require specific experimental conditions (Liu et al., 2019). Therefore, in this study, a k-means clustering method was employed to automatically classify and organize all samples based on their gene expression patterns, forming distinct clusters. By annotating these clusters, each cluster can represent specific conditions.

Based on the comparative analysis from three perspectives, we observed significant differences between the global module and the cluster-specific modules in terms of both gene composition and the distribution of biological functions. Furthermore, compared to the co-expression modules obtained from the global samples, the co-expression modules corresponding to the clustered samples indeed showed a significant improvement in accuracy. Specifically, under specific conditions, there was a higher similarity between the gene expression profiles and the average expression profiles of the modules they belonged to. Additionally, we found that the Cluster 1 specific module has the most unique gene composition and the most concentrated biological functions compared to the global module, which warrants further investigation.

Although the co-expression analysis of the clustered samples has higher accuracy, it does not mean that the analysis of the global samples becomes meaningless. On the contrary, by combining the co-expression analysis of the clustered samples with that of the global samples, we can obtain more valuable information. In this study, a correlation difference value (CDV) was proposed to explain the condition specificity of a gene by comparing the correlation between the gene expression profile and the average expression profile of the cluster-specific module, and the correlation between the gene expression profile and the average expression profile of the global module under specific conditions. CDV has been demonstrated in this paper to reflect the impact of a gene on the similarity between the cluster-specific module and the global module. Genes with higher CDV values exhibit higher condition specificity and are worth further investigation.

In this study, through the investigation of the specific condition Cluster1, a gene with high condition specificity and high betweenness centrality in the biological function “PPR-type RNA editing factor activities” was identified, namely CSS0047732.1. CSS0047732.1 is a PPR-type RNA editing factor (CWM1), and its expression profile confirms that it exhibits high expression in the bud leaves of “Anji Baicha” during the bud-prealbinism stage. This indicates that the gene CSS0047732.1 plays an important role during the bud-prealbinism stage in “Anji Baicha”.

Previously, researchers have found that the expression level of a PPR gene significantly increases during the albino stage of “Anji Baicha”, and PPR genes are involved in the albino phenotypes of many plants (Yuan et al., 2015). The correlation between PPR genes and albino phenotypes has been observed in several plants (Yu et al., 2009; Lan et al., 2010; Williams and Barkan, 2003; Gothandam et al., 2005; Cui et al., 2019). Previous studies have shown that undeveloped plastids without obvious internal membrane structures are present in the albino leaves of “Anji Baicha”, indicating that chloroplast development in “Anji Baicha” is suppressed (Li et al., 2011). Therefore, CSS0047732.1, as a PPR-type RNA editing factor (CWM1), may be upregulated and involved in the process of inhibiting chloroplast development, leading to albino phenotypes in “Anji Baicha”.

## Supporting information

Figure S1. Pseudoaligned reads percentage and sequencing reads distribution of the Camellia sinensis RNA-Seq samples.

Figure S2. Metadata enrichment heatmap of k-means clusters.

Figure S3. Scale-free topology fitting index plot and mean connectivity plot of weighted gene co-expression network analysis.

Figure S4. Hierarchical clustering dendrogram of module eigengenes.

Figure S5. Hierarchical clustering dendrogram and module coding color of weighted gene co-expression network analysis.

Figure S6. Correlation analysis between cluster1-specific co-expression modules and the global co-expression module.

Figure S7. Functional enrichment heatmap of cluster1-specific co-expression modules and the global co-expression module.

Figure S8. Illustrative graph demonstrating the change in module similarity as the threshold increases from 0.3 to 0.6.

Table S1. Annotation information for RNA-Seq samples.

Table S2. Expression profile with 50,525 genes as columns and 769 RNA-Seq samples as rows, with TPM values used as expression levels.

Table S3. Annotation information for Camellia sinensis genes.

Table S4. Enrichment analysis of metadata terms for cluster samples.

Table S5. Module coding color of Camellia sinensis genes, and the GMC and CDV of each gene.

Table S6. Association analysis between the global module and cluster-specific modules.

Table S7. Functional enrichment analysis of module genes using Mapman annotations as biological functions.

Table S8. Network analysis of functional genes in co-expression modules.

## Acknowledgments

X.Z. is sponsored by a China Scholarship Council fellowship.

## Author contributions

X.Z. led the main work of this study, including project conception, data annotation, data analysis, and paper writing. P.K.L. contributed suggestions on using K-means clustering analysis to address sample imbalance issues in co-expression analysis. M.M. and Y.W. co-supervised X.Z. in completing this project. M.M. and P.K.L. both participated in the revision of the paper. The authors thank all members of the Mutwil Lab for their suggestions and assistance with this manuscript.

## Supplementary tables

**Table S1. Annotation information for RNA-Seq samples, including cultivar, tissue, sampling age, experimental treatment, pseudoaligned rate, and k-means cluster IDs under different n_clusters.**

**Table S2. Expression profile with 50,525 genes as columns and 769 RNA-Seq samples as rows, with TPM values used as expression levels.**

**Table S3. Annotation information for *Camellia sinensis* genes, including ITAK annotation, KEGG annotation, GO annotation, Pfam annotation, and Mapman annotation.**

**Table S4. Enrichment analysis of metadata terms for cluster samples.** (A-C) FDR values for cultivars, tissues and experimental treatments, respectively. (D-F) Rich factors for cultivars, tissues and experimental treatments, respectively.

**Table S5. Module coding color of *Camellia sinensis* genes in the global module and cluster-specific modules, and the gene-module consistency coefficient (GMC) and correlation difference value (CDV) of each gene.** (A) Module coding colors. (B-H) gene-module consistency coefficient (GMC) and correlation difference value (CDV) of each gene in Cluster1-Cluster7, respectively.

**Table S6. Association analysis between the global module and cluster-specific modules.** (A-G) FDR values for Cluster1-Cluster7, respectively. (H-N) Jaccard index for Cluster1-Cluster7, respectively.

**Table S7. Functional enrichment analysis of module genes using Mapman annotations as biological functions.** (A-H) FDR values for the global and Cluster1-Cluster7 samples, respectively. (I-P) Rich factors for the global and Cluster1-Cluster7 samples, respectively.

**Table S8. Network analysis of functional genes in co-expression modules.** (A-E) Co-expression networks related to the R2R3-MYB transcription factor family (cluster1-darkorange2, cluster1-white, global-darkmagenta, global-darkolivegreen, and global-royalblue, respectively). (F-M) Co-expression networks related to Photosystem II (cluster1-darkturquoise, cluster1-lightcyan, cluster1-orangered4, cluster1-steelblue, global-brown, global-mediumorchid, global-salmon4, and global-yellow, respectively). (N-R) Co-expression networks related to PPR-type RNA editing factor activities (cluster1-grey, cluster1-lightsteelblue1, cluster1-yellow4, global-cyan, and global-yellow, respectively). (S-X) Co-expression networks related to RING E3 ubiquitin ligase activities.RING-H2-class (cluster1-brown, cluster1-cyan, global-darkmagenta, global-pink, global-saddlebrown, and global-tan, respectively).

## Supplementary figure

**Figure S1.** P**seudoaligned reads percentage and sequencing reads distribution of the *Camellia sinensis* RNA-Seq samples.**

**Figure S2. Metadata enrichment heatmap of k-means clusters.** Orange/gray dots distinguish significant/non-significant enrichments, and the size of the dots represents the rich factor. (A) Cultivar. (B) Tissue. (C) Experimental treatment.

**Figure S3. Scale-free topology fitting index plot and mean connectivity plot of weighted gene co-expression network analysis.** (A-H) shows Cluster 1-Cluster 7 and the global group, respectively.

**Figure S4. Hierarchical clustering dendrogram of module eigengenes.**(A-H) shows Cluster 1-Cluster 7 and the global group, respectively.

**Figure S5. Hierarchical clustering dendrogram and module coding color of weighted gene co-expression network analysis.** (A-H) shows Cluster 1-Cluster 7 and the global group, respectively.

**Figure S6. Correlation analysis between cluster1-specific co-expression modules and the global co-expression module.** The presence or absence of asterisks (*) in the cells distinguishes significant or non-significant enrichment, and the color intensity of the cells represents the Jaccard index. The bar chart displays the size of the gene sets.

**Figure S7. Functional enrichment heatmap of cluster1-specific co-expression modules and the global co-expression module.** The presence or absence of asterisks (*) in the cells indicates significant or non-significant enrichment, and the color of the cells represents the rich factor. The bipartite graph in the central region of the figure reveals the relationship between cluster-specific co-expression modules and the global co-expression module. In the graph, nodes represent modules, and the color intensity of nodes represents their degree. The presence or absence of connecting lines between nodes indicates significant or non-significant associations, and the color intensity of the connecting lines represents the Jaccard index.

**Figure S8. Illustrative graph demonstrating the change in module similarity as the threshold increases from 0.3 to 0.6.**

