## Supplementary figures and images for "A method for mining condition-specific co-expressed genes in *Camellia sinensis* based on K-means clustering: A case study of “Anji Baicha” tea cultivar"

### Figure S1. Pseudoaligned reads percentage and sequencing reads distribution of the Camellia sinensis RNA-Seq samples.

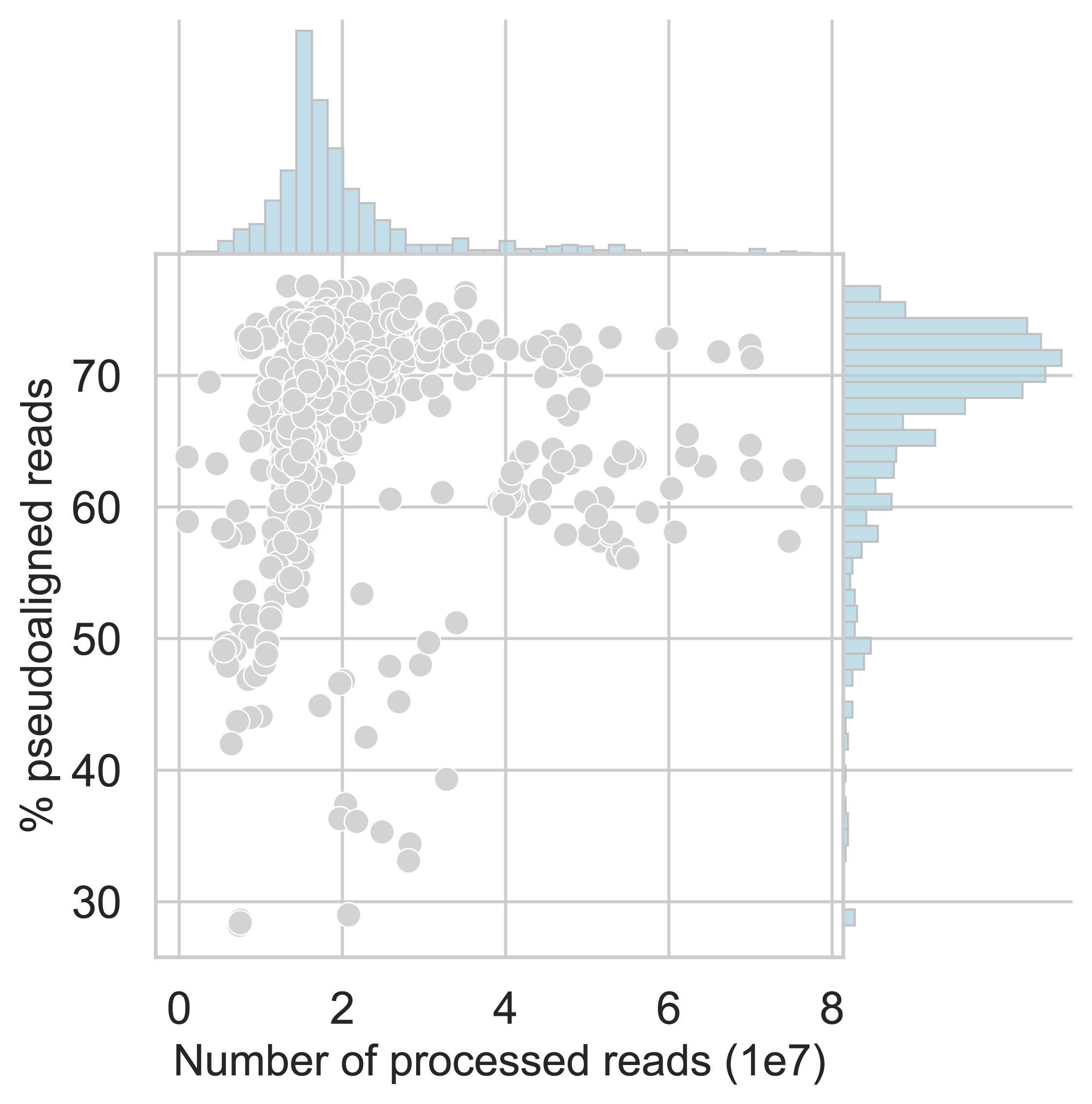

### Figure S2. Metadata enrichment heatmap of k-means clusters.

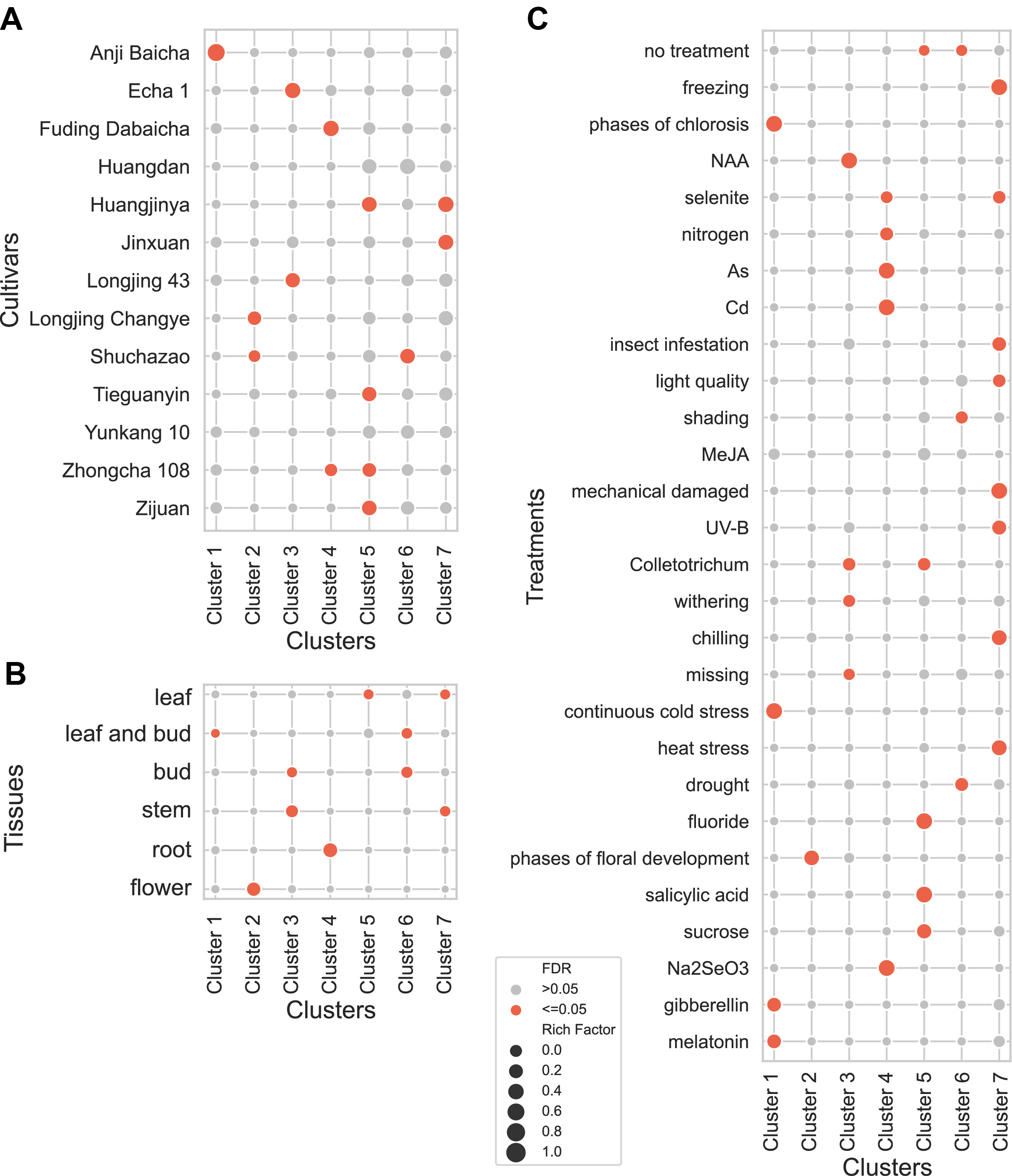

### Figure S3. Scale-free topology fitting index plot and mean connectivity plot of weighted gene co-expression network analysis.

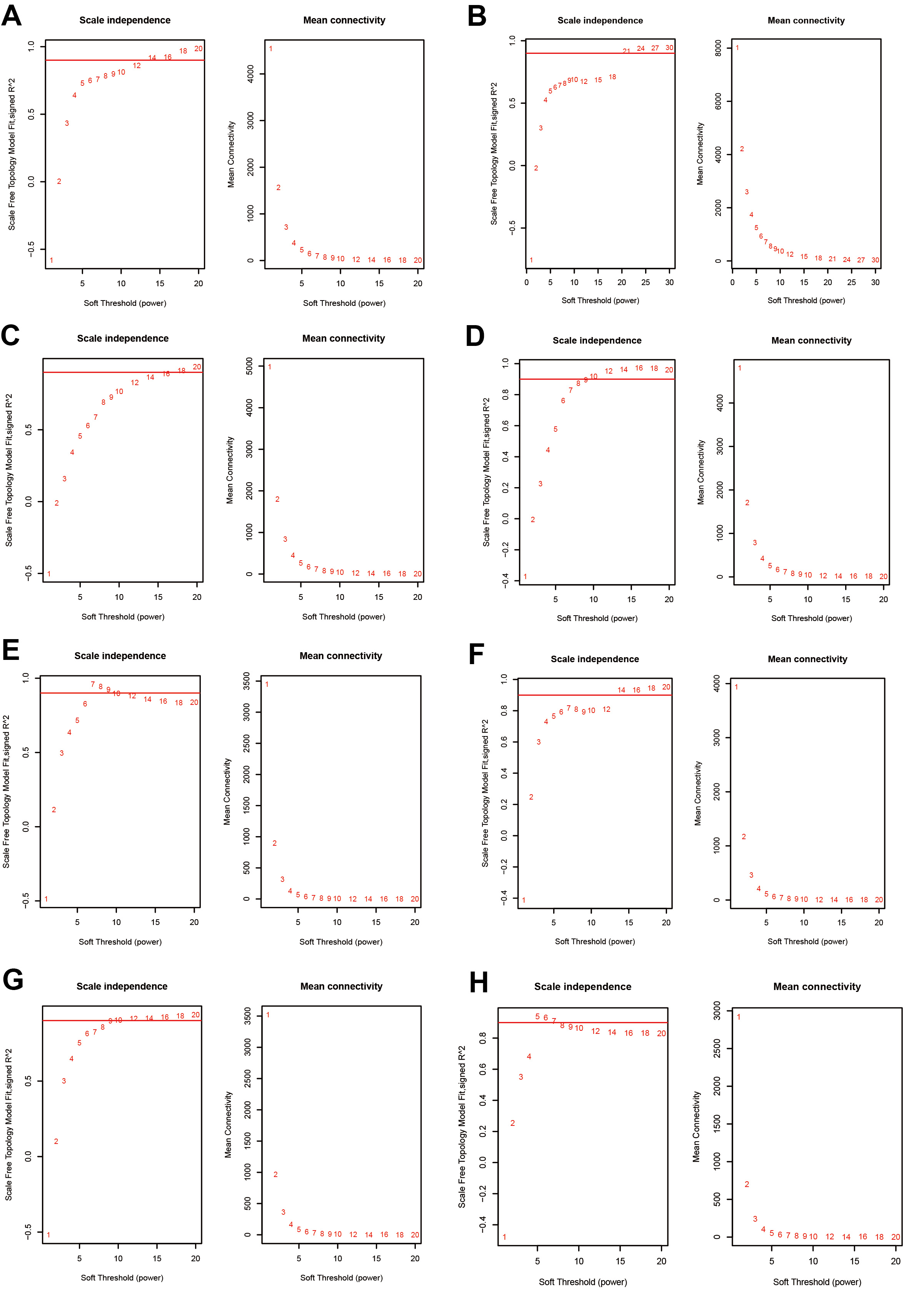

### Figure S4. Hierarchical clustering dendrogram of module eigengenes.

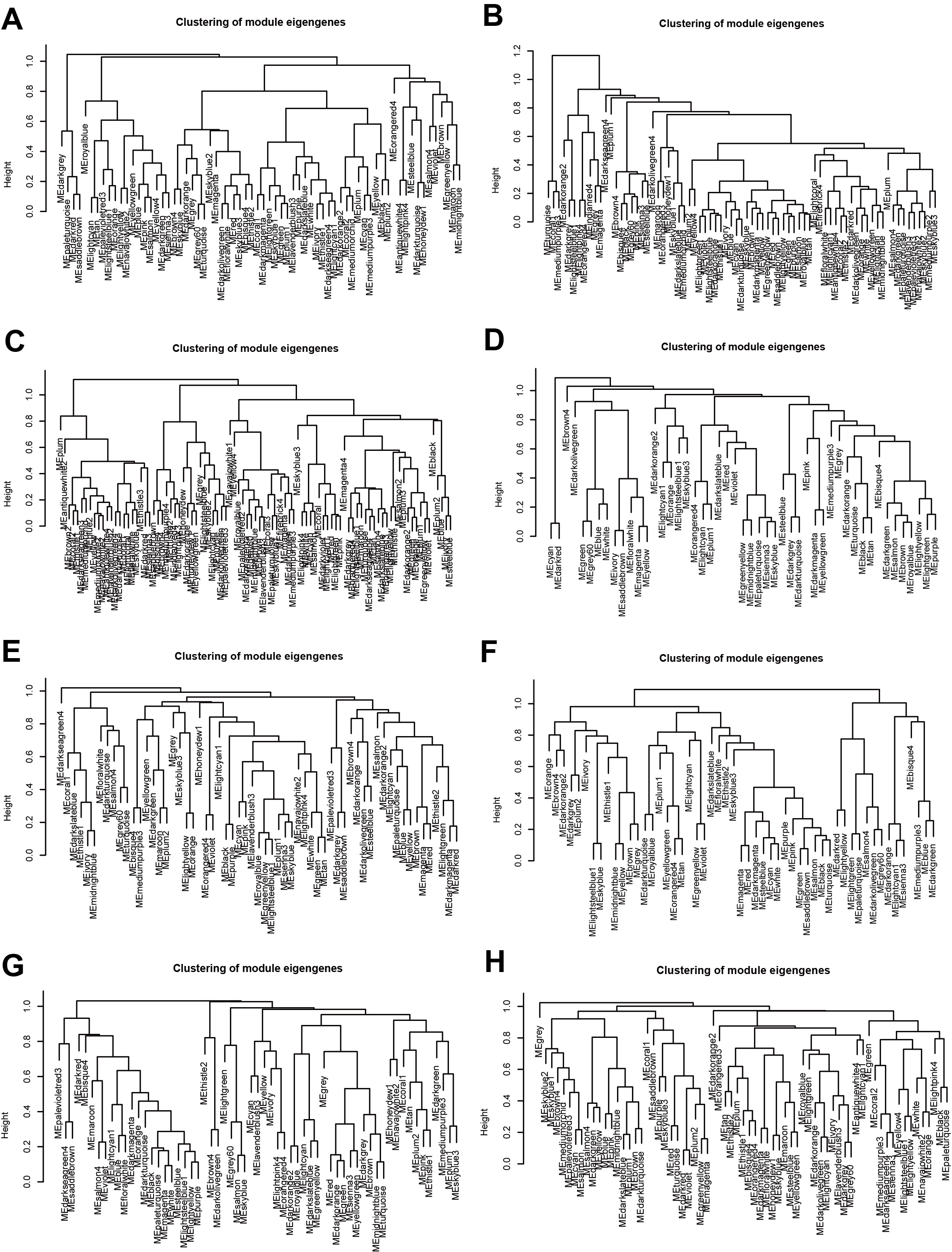

### Figure S5. Hierarchical clustering dendrogram and module coding color of weighted gene co-expression network analysis.

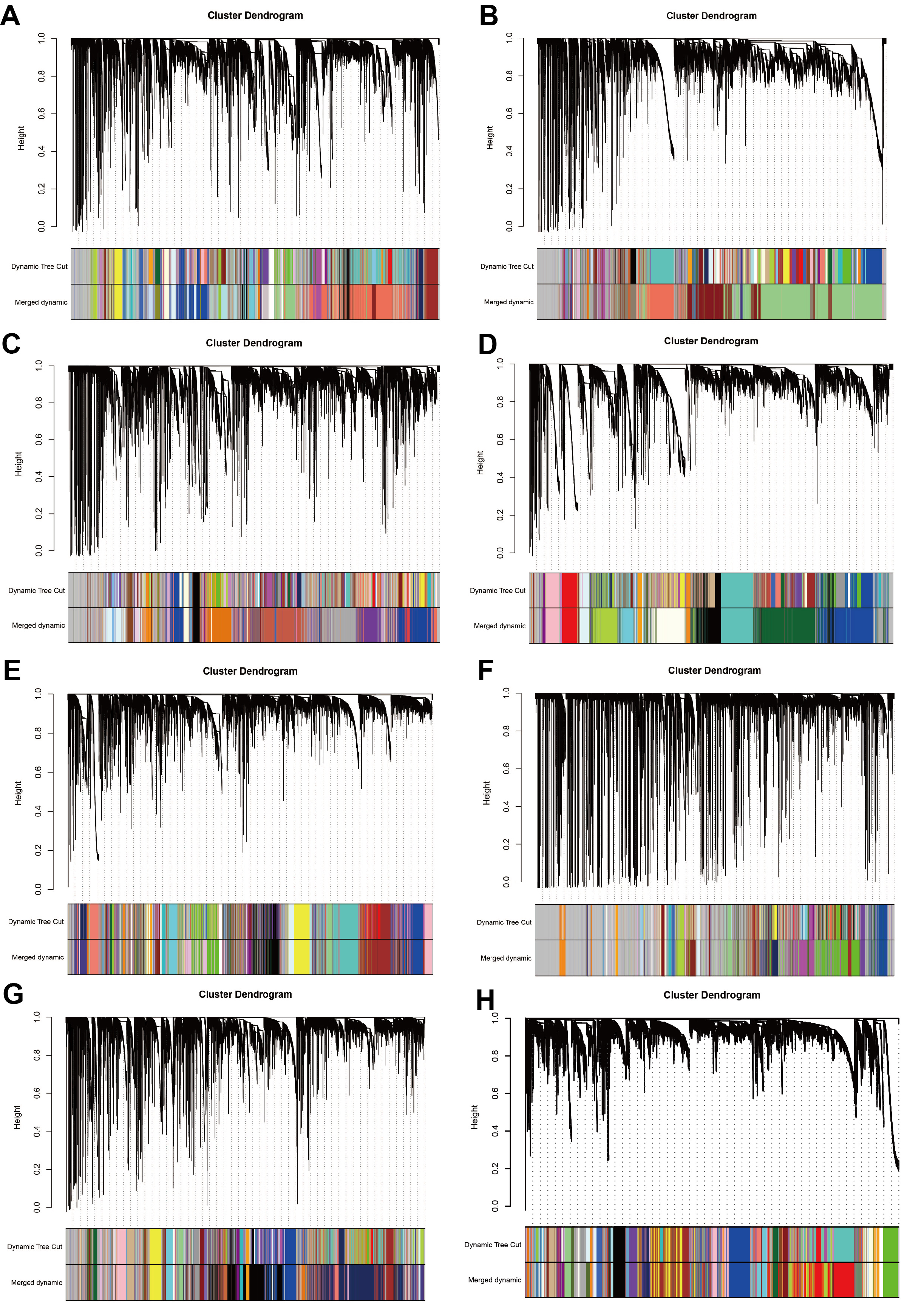

### Figure S6. Correlation analysis between cluster1-specific co-expression modules and the global co-expression module.

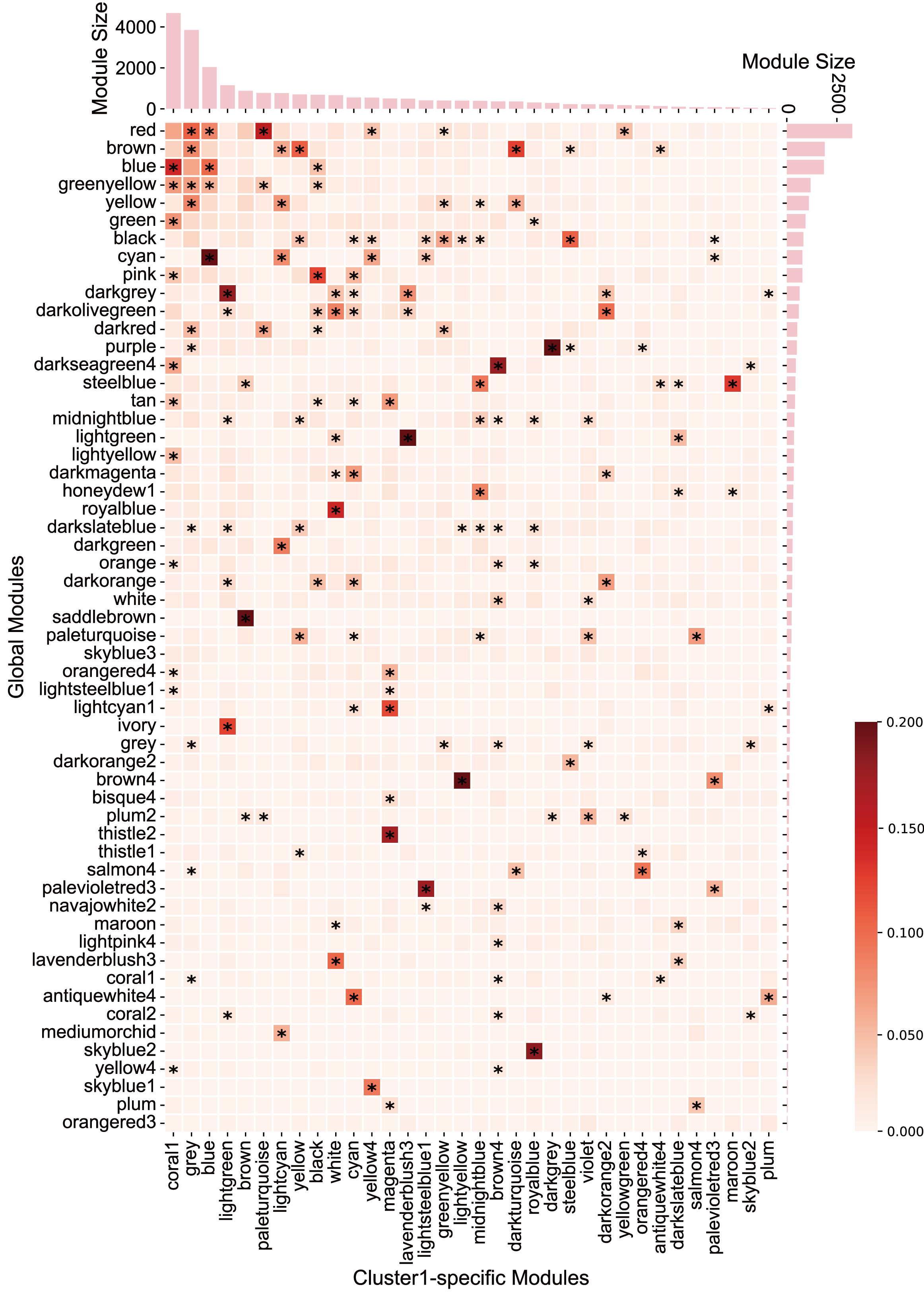

### Figure S7. Functional enrichment heatmap of cluster1-specific co-expression modules and the global co-expression module.

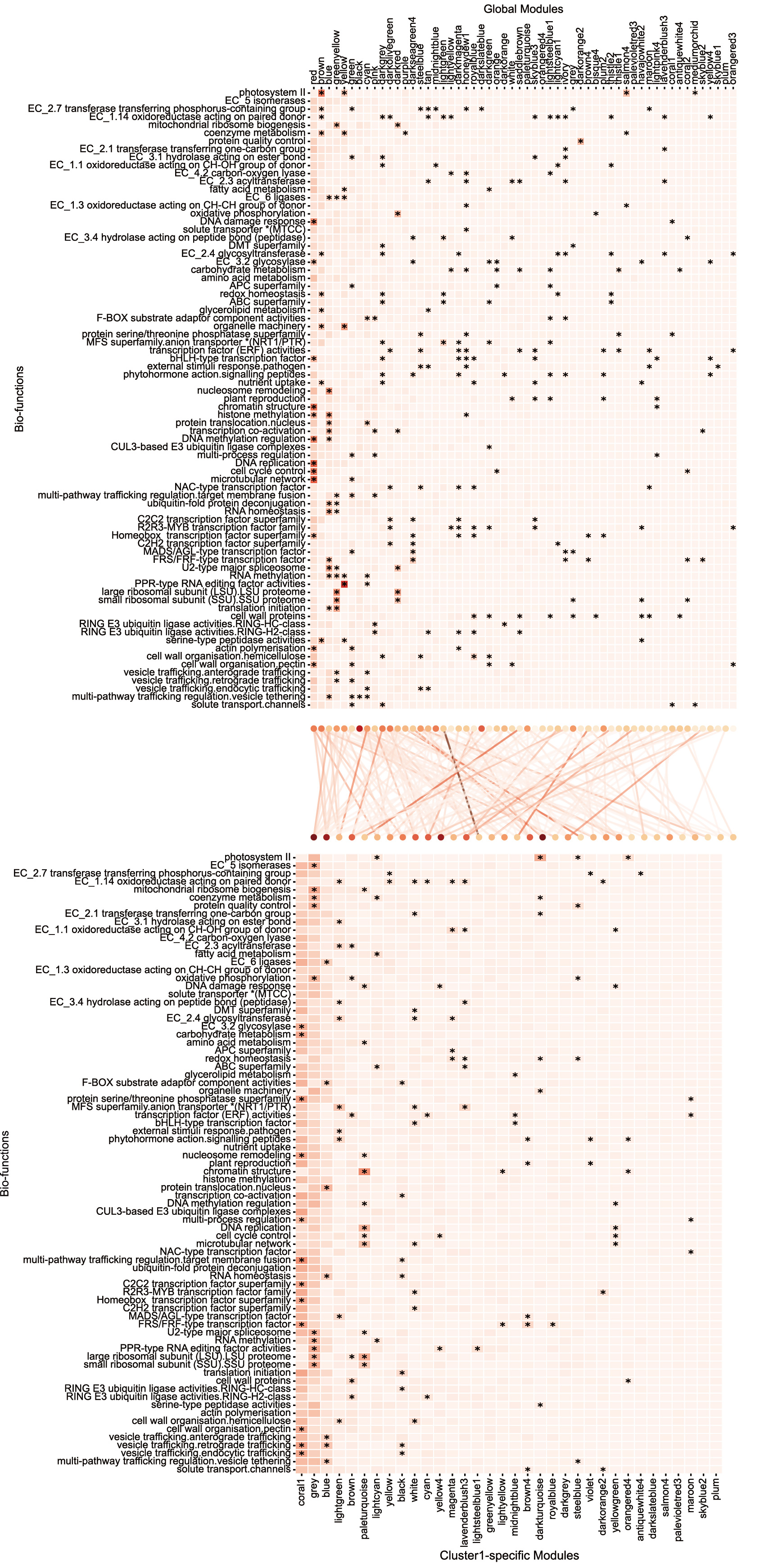

### Figure S8. Illustrative graph demonstrating the change in module similarity as the threshold increases from 0.3 to 0.6.

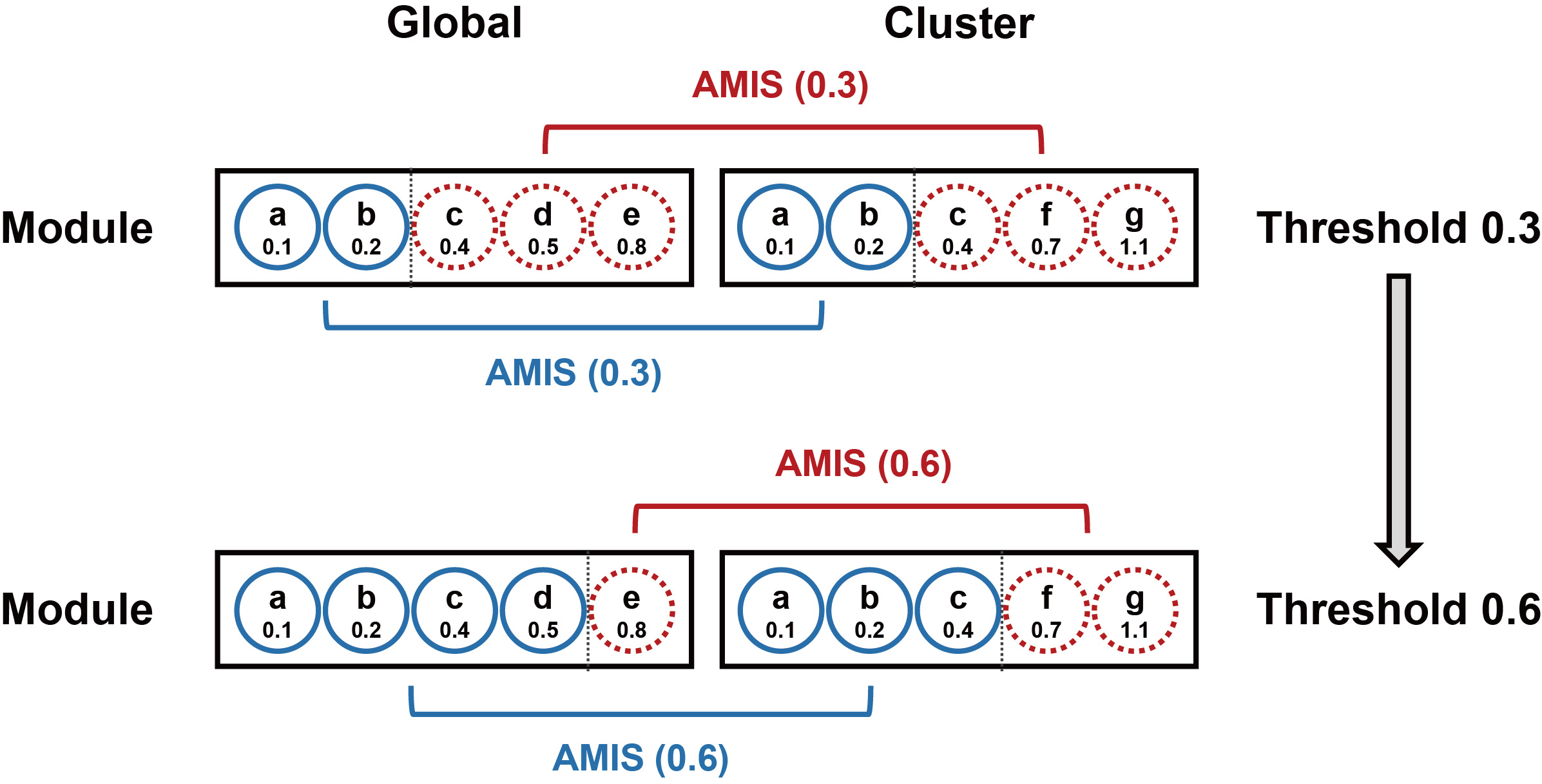
